# OrgaMapper: A robust and easy-to-use workflow for analyzing organelle positioning

**DOI:** 10.1101/2023.07.10.548452

**Authors:** Christopher Schmied, Michael Ebner, Paula Samsó Ferré, Volker Haucke, Martin Lehmann

**Author notes:** equal contribution (Christopher Schmied, Michael Ebner).

## Abstract

Eukaryotic cells are highly compartmentalized by a variety of organelles that carry out specific cellular processes. The position of these organelles within the cell is elaborately regulated and vital for their function. For instance, the position of lysosomes relative to the nucleus controls their degradative capacity and is altered in pathophysiological conditions. The molecular components orchestrating the precise localization of organelles remain incompletely understood. A confounding factor in these studies is the fact that organelle positioning is surprisingly non-trivial to address. E.g., perturbations that affect the localization of organelles often lead to secondary phenotypes such as changes in cell or organelle size. These phenotypes could potentially mask effects or lead to the identification of false positive hits. To uncover and test potential molecular components at scale, accurate and easy to use analysis tools are required that allow robust measurements of organelle positioning.

Here, we present an analysis workflow for the faithful, robust, and quantitative analysis of organelle positioning phenotypes. Our workflow consists of an easy to use Fiji plugin and an R Shiny App. These tools enable users without background in image or data analysis to (1) segment single cells and nuclei and to detect organelles, (2) to measure cell size and the distance between detected organelles and the nucleus, (3) to measure intensities in the organelle channel plus one additional channel, and (4) to plot the results in informative graphs. Using simulated data and immunofluorescent images of cells in which the function of known factors for lysosome positioning has been perturbed, we show that the workflow is robust against common problems for the accurate assessment of organelle positioning such as changes of cell shape and size, organelle size and background.

## Introduction

Eukaryotic cells are highly compartmentalized by membrane-enclosed and membrane-less organelles. Compartmentalization of the cytoplasm into the semi-confined spaces of specialized organelles ensures the efficient channeling of all fundamental biological processes that are hallmarks of eukaryotic life. The position of these organelles within the cytoplasm is highly specific, subject to regulation by metabolic cues and signaling pathways, and interdependent with organelle identity and function (van Bergeijk, Hoogenraad et al. 2016, Barlan and Gelfand 2017). The wide range of human diseases that are associated with organelle transport is testimony to the paramount importance of the fidelity of organelle positioning (Aridor and Hannan 2000, Mandelkow and Mandelkow 2002, van Bergeijk, Hoogenraad et al. 2016). The transport and dynamic positioning of organelles is governed by specialized molecular motors as most dramatically exemplified in highly polarized cells such as neurons (Hirokawa, Niwa et al. 2010). These molecular motors use the actin and microtubule cytoskeleton as tracks and a plethora of scaffolding, regulatory, and signaling proteins to connect to specific cargo organelles. More recently, membrane contact sites between different organelle species were uncovered as an additional layer in the complex regulatory network that coordinates organelle positioning (Neefjes, Jongsma et al. 2017)

Molecular mechanisms and functional relevance of organelle positioning are the focus of intense research with examples ranging from early endosomes (Caviston, Zajac et al. 2011, Ketel, Krauss et al. 2016) recycling endosomes (Winter, Hopfner et al. 2012) autophagosomes (Kimura, Noda et al. 2008, Wijdeven, Janssen et al. 2016), peroxisomes (Baker, Sparkes et al. 2010, Smith and Aitchison 2013), lipid droplets (Thiam and Beller 2017), mitochondria (Okatsu, Saisho et al. 2010, Schuler, Lewandowska et al. 2017, Kraft and Lackner 2018), to the Golgi complex (Gurel, Hatch et al. 2014, Song, Gras et al. 2019). Notably, in the last decade the positioning of lysosomes and late endosomes has specifically gained attention as it is inextricably linked to metabolic homeostasis and growth signaling (Bonifacino and Neefjes 2017, Cabukusta and Neefjes 2018). Even the intracellular distribution and function of membrane-less organelles such as stress granules is under regulation by microtubule-based transport (Ivanov, Chudinova et al. 2003, Loschi, Leishman et al. 2009). In patient derived cells, altered organelle positioning can hint at possible molecular underpinnings of disease or provide cues regarding putative disease mechanisms (Huizing, Hess et al. 2004, Wali, Sutharsan et al. 2016).

Evidently, there is a recent surge in efforts to decipher the physiological cues and molecular components that govern organelle positioning in health and disease and many novel cell biological concepts and disease links were uncovered on the way. However, our understanding of these processes is far from complete. For instance, it is mostly unclear which factors on organelle membranes bridge to the molecular motor machinery, how and via which factors organelles signal to the machinery, and how the transport machinery is instructed by external cues. In order to identify factors involved in these mechanisms and to facilitate the use of organelle positioning as a diagnostic readout (Huizing, Hess et al. 2004), biologists and pathologists require easy to use and scalable tools. These tools must be able to faithfully and robustly localize organelles in images and provide easily interpretable measurements and analysis.

There are many general-purpose solutions and platforms available, such as Fiji, Icy, Cell Profiler, Knime, or Illastik, that enable scientists to implement image analysis workflows (de Chaumont, Dallongeville et al. 2012, Schindelin, Arganda-Carreras et al. 2012, Berg, Kutra et al. 2019, Dietz, Rueden et al. 2020, Stirling, Swain-Bowden et al. 2021). However, creating a robust and easy to use analysis workflow for complex biological phenomena such as organelle positioning requires specialized expertise in image analysis and software engineering. To allow biologists with little to no background in programming and data science to execute advanced image and data analysis, well-tested workflow templates are required that use established components, are provided to the user with comfortable graphical user interfaces (GUIs), and are supported with ample documentation (Miura and Norrelykke 2021). Fortunately, this need has become recognized in the scientific community, creating the emerging field of bioimage analysis (Cimini, Norrelykke et al. 2020) with such workflow templates being increasingly published and made available to the broader community (Erguvan, Louveaux et al. 2019, Klickstein, Mukkavalli et al. 2020, Fisch, Evans et al. 2021, Schmied, Soykan et al. 2021).

We reviewed published approaches to quantify organelle positioning and encountered numerous different strategies, most of them based on manual or semi-automatic custom-made solutions either using intensity or the distance of individual objects as measurement read out (**Supplementary table 1**). Here, we have addressed advantages and shortcomings of different basic analysis strategies using simulated data. We found that organelle positioning is surprisingly non-trivial to address as it is highly sensitive to cell size, cell shape, organelle shape, background intensity or organelle intensity distributions. Based on these findings we have implemented detection- and intensity-based analysis in a complete open source image and data analysis workflow as an ImageJ/Fiji Plugin and an easy-to-use R Shiny App called OrgaMapper. This workflow enables users with little background in image and data analysis to robustly quantify organelle positioning while accounting for compounding factors in the datasets, such as differences in cell size or organelle shape. We validated the analysis workflow using immunofluorescence and live-cell imaging datasets. The entire workflow, accompanying test data and extensive documentation are open source and freely available to the community. We thus provide the rapidly expanding field of organelle positioning with a publicly accessible, validated, fully automated, and easy to use analysis workflow.

## Availability and support

The image analysis workflow is available as a Fiji plugin and can be installed via an update site: https://sites.imagej.net/Cellular-Imaging/. The source code of the R Shiny App can be downloaded from github: https://github.com/schmiedc/OrgaMapper_Rshiny. Documentation for the usage of the Fiji plugin and R Shiny App can be found here: https://schmiedc.github.io/OrgaMapper/. The software, test datasets and settings files can be found in the following zenodo repository: https://doi.org/10.5281/zenodo.8128670

### Contact and support

https://forum.image.sc/u/schmiedc/.

## Results

### Distance of individual organelles from the nucleus is a robust readout of organelle position

In order to untangle the complex molecular machinery that governs the positioning of organelles, high content and high throughput microscopy screening can be used to test many different molecular components at scale using pharmacological and genetic treatments. When using light microscopy images there are two major means of assessing the relative position of organelles in the cytoplasm: First, by plotting the intensity of a fluorophore which decorates the organelle in relation to a point of reference (**Figure 1A**) (Rocha, Kuijl et al. 2009, Li, Rydzewski et al. 2016, Starling, Yip et al. 2016, Filipek, de Araujo et al. 2017, Hong, Pedersen et al. 2017, Willett, Martina et al. 2017, Walton, Patel et al. 2018, Tapia, Jimenez et al. 2019)(**Supplementary table 1**). Second, by detecting or segmenting the organelles and measuring the distance of individual objects to a point of reference (**Figure 1B**) (Marat, Wallroth et al. 2017, Ba, Raghavan et al. 2018, Klickstein, Mukkavalli et al. 2020)(**Supplementary table 1**). In order to test advantages and disadvantages of these two approaches, we used images of idealized simulated cells consisting of nucleus, whole cell, and vesicles (**Figure 1C**). We measured the integrated density of vesicles in the total cell (I_t_) and within a defined perimeter equidistant to the nucleus (I_p_) and calculated the ratio (I_p_/I_t_), also known as the perinuclear index (**Figure 1A**) (Li, Rydzewski et al. 2016, Willett, Martina et al. 2017, Tapia, Jimenez et al. 2019) as well as the actual distance of the simulated vesicles from the nuclear boundary (**Figure 1B**). We then computed the measurement error that the variation of defined morphological parameters would introduce for both methods and dubbed it the *error factor* (see methods for detailed description). The larger the *error factor* the more sensitive the method is to the morphological parameter.

**Fig. 1:**
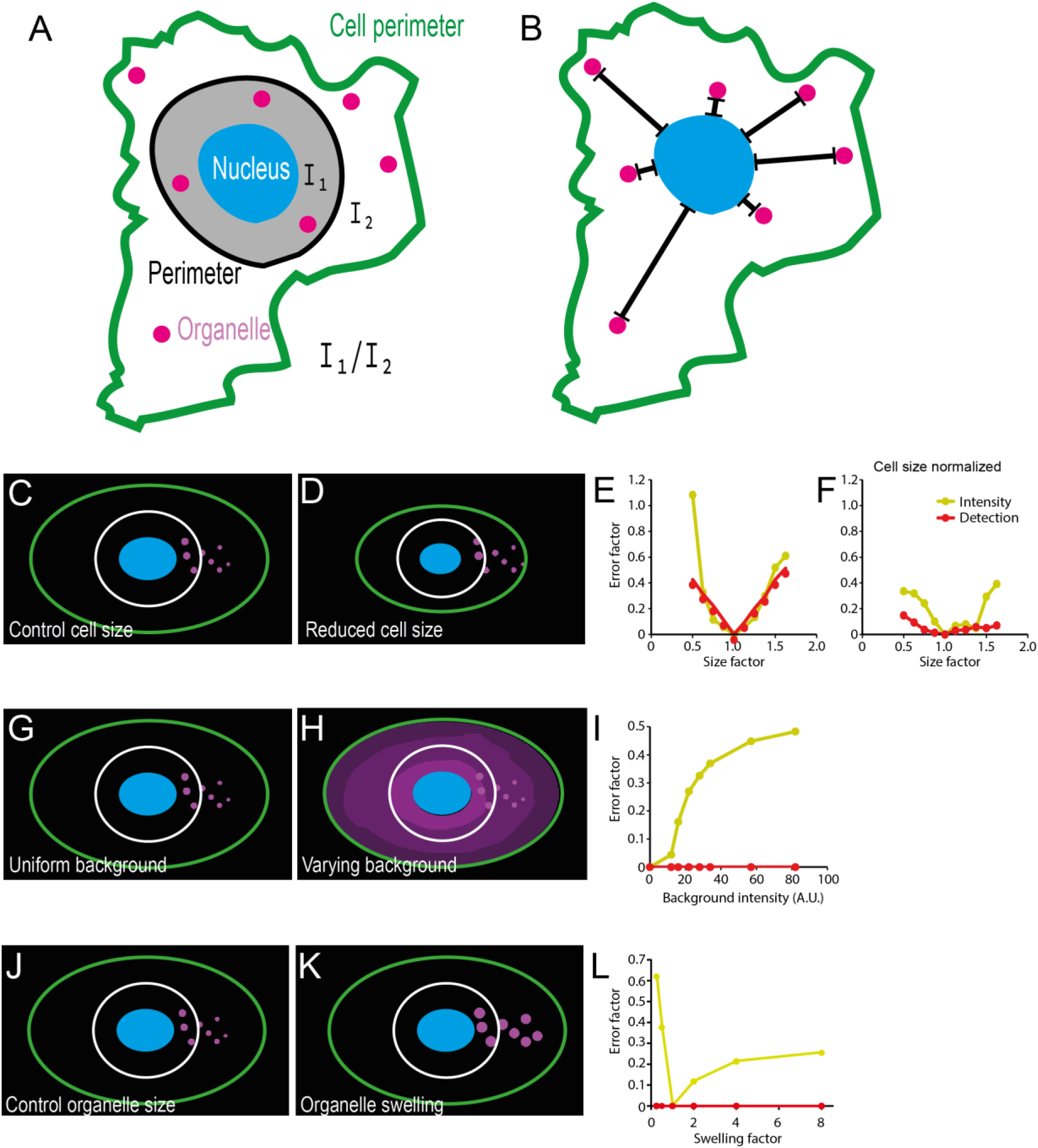
Measuring the distance of individual organelles from the nucleus is robust to changes in cell and organelle morphology *in silico*. Principle of analyzing organelle position using intensity, i.e. perinuclear index; I_1_…intensity within a perimeter equidistant to the nucleus, I_2_…intensity outside of the perimeter (**A**) or via measuring the distance of individually detected lysosomes from the nucleus (**B**). (**C**) Simulated cell for testing robustness of organelle positioning measurements (blue: nucleus, magenta: organelles, white: perimeter for intensity ratio, green: cell boundary). (**D-F**) Measuring the distance of individual organelles is robust to changes in cell size when normalized by Feret’s diameter. (**G-I**) Measurement of distance of individual organelles is robust to fluctuations in background fluorescence. (**J-L**) Measuring the distance of individual organelles is robust to changes in organelle size.

The positioning of late endomes and lysosomes (hereafter referred to as lysosomes for simplicity) directly affects cell growth and therefore cell size (Korolchuk, Saiki et al. 2011). We reasoned that altered cell size would skew the results of organelle positioning measurements. Indeed, using our simulated cells we found that altered cell size could introduce a substantial error when performing distance measurements and especially when performing intensity-based quantification (**Figure 1C-E**). Normalizing intensity with measurement area decreased the error, suggesting that factoring in cell size could at least partially prevent systematic bias when organelle positioning is quantified by intensity-based methods. Normalizing distance by the cell’s diameter (Feret’s diameter) almost completely prevented the error introduced by the altered cell size (**Figure 1F**). Further, basing positioning on intensity measurements is only feasible under the assumption that there is a linear relationship between the intensity of the fluorophore and the occurrence of the organelle. However, uneven distribution of background signal due to inherently low z-resolution in light microscopy as well as staining artifacts could alter the result erroneously (**Figure 1G-I**). In contrast, when using distance measurement of individual organelles, the influence of the background signal and possible staining artifacts may be eliminated (**Figure 1I**). Additionally, organelle size (**Figure 1J-L**) or organelle intensity changes due to an experimental treatment cannot be accounted for in intensity-based analysis (e.g. interference with endo-lysosome homeostasis frequently results in vesicle swelling or altered acidity, which would affect the intensity of frequently used endo-lysosomal markers such as lysotracker dyes). Finally, only by detecting individual organelles potentially different subpopulations of organelles can be identified and analyzed further.

Together, our modeling approach suggests that segmenting individual organelles combined with cell size normalization is the most robust and faithful quantification method for organelle positioning. Based on these results we developed an ImageJ plugin with adjustable parameters for detection and distance measurements of individual organelles in single cells.

### Interactive segmentation and detection using OrgaMapper

In typical high throughput imaging approaches, different cellular components such as the nucleus, cytoplasm and the cellular compartment of interest are labeled with appropriate stains or fluorescent markers that report cellular and intracellular architecture (**Figure 2A**). To assess the resulting phenotype of the treatment, the labeled structures need to be segmented and measured automatically in a robust manner.

**Fig. 2:**
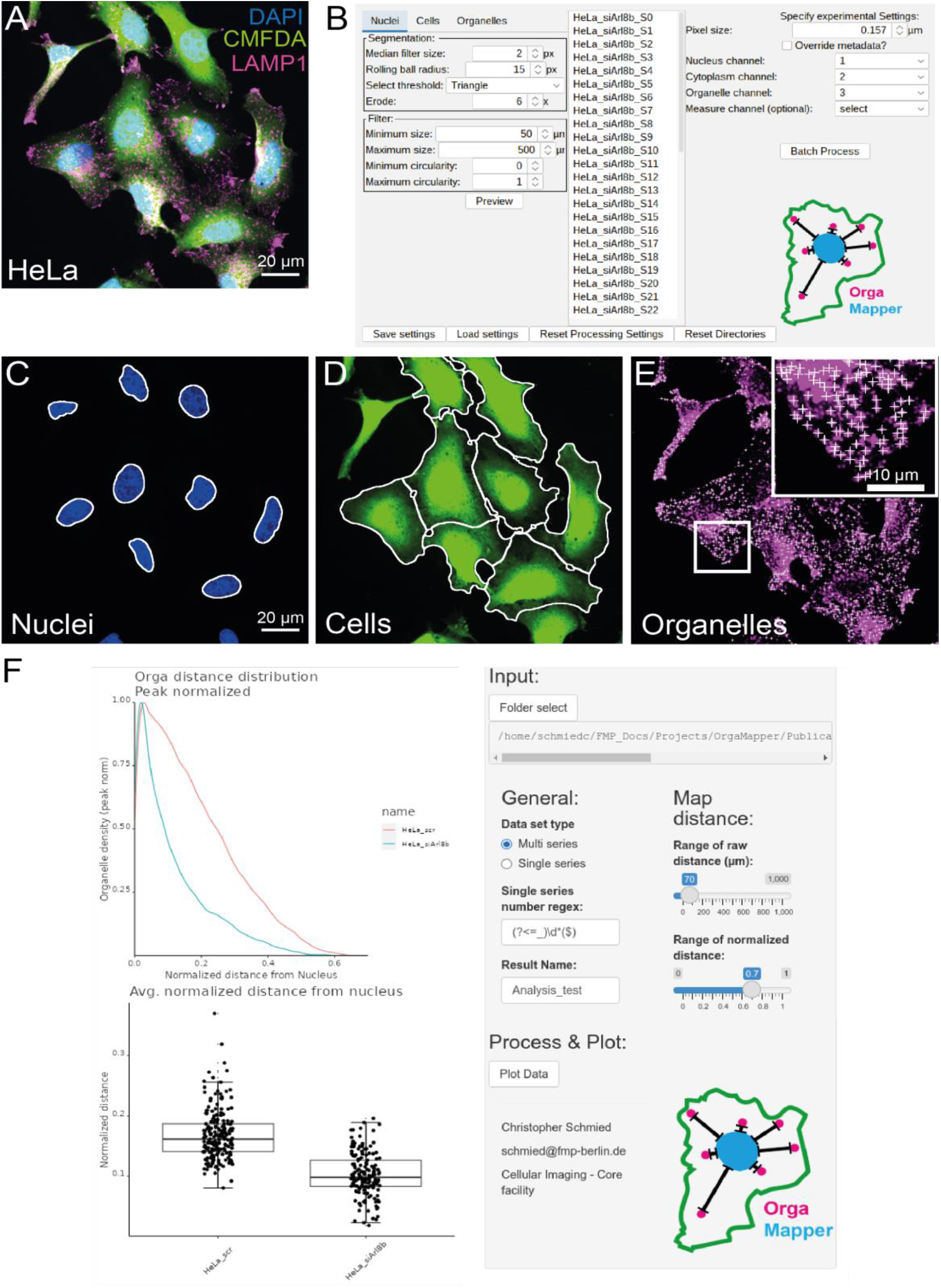
OrgaMapper enables interactive adjustment of segmentation and detection. (**A**) Typical input data with labels for nuclei (DAPI), cytoplasm (CMFDA) and organelle channel (LAMP1, i.e. lysosomal marker). (**B**) The user can adjust and test the segmentation and detection parameters in the GUI of OrgaMapper on all the available files and execute the image analysis in batch. (**C**) Segmentation of the nucleus. (**D**) Cell segmentation filtered for segmented nuclei. (**E**) Organelle detection excluding organelles overlapping with the nucleus. (**F**) OrgaMapper R Shiny graphical user interface.

OrgaMapper implements the necessary image processing and analysis tasks in one convenient ImageJ plugin. The segmentation can be adjusted via a graphical user interface (GUI) that allows users to easily and interactively adjust and review the segmentation and detection over the entire dataset and execute the image analysis in batch (**Figure 2B**). For reviewing the performance of the image analysis the segmentation and detection ROIs are provided as overlays over the different image channels (**Figure 2C-E**). All the necessary analysis parameters are documented and saved in a settings file that can be loaded by the plugin for reproduction. The nuclei are segmented using an automatic intensity-based threshold (**Figure 2C**). Nuclei at the edge of the field of view, as well as nuclei that are too small, large or with unsuitable shape are rejected. For the segmentation of the cell area a marker-controlled watershed is employed to segment and separate the cells. Cells without a valid nucleus segmentation or with multiple nuclei are rejected from further analysis (**Figure 2D**). In order to robustly detect the peaks of individual organelles within a certain size range an implementation of the Laplacian of Gaussian (LoG) (Sage, Neumann et al. 2005) is applied and subsequently the maxima are detected (**Figure 2E**).

The segmentation masks of individual nuclei and cells are used to compute for each individual pixel in the cytoplasm the shortest distance to the closest nucleus pixel (i.e. Euclidean distance). This yields a Euclidean distance map (EDM) of the cytoplasm with respect to the edge of the nucleus. The organelle detections are then applied to the EDM within the cytoplasm to extract the location of each organelle in relation to the edge of the nucleus resulting in the generation of a detection map. Further, the intensity is measured at the detection localization in the organelle channel. As a further parameter independent from the detection of the organelles, for each pixel in the cytoplasm the gray value in the organelle channel is extracted together with the distance based on the EDM to generate an intensity map. The analysis further extracts per-cell key morphological parameters such as diameter and area as well as overall statistics such as the number of detections as well as the average intensity in the organelle channel within the entire segmented cytoplasm. To estimate a background value, the mean intensity per field of view outside of the unfiltered cell segmentation is measured (**Supplementary Table 3**). Finally, individual nuclei, cell and detection ROIs are saved and made accessible for further analysis.

After execution of the Fiji plugin the analysis results can be loaded into the provided R Shiny App, which automatically collects and saves all measurements in a summary table in the specified output directory. A basic analysis with plots based on the measured parameters is provided by default and visualized in the plotting interface of the R Shiny App. Key settings for the data processing (i.e. background subtraction, cell diameter filter) and plotting (e.g. plotting ranges and bin width) can be adjusted via a GUI (**Figure 2F**).

### Spatial statistical analysis enables accurate measurement of organelle positioning phenotypes

We validated the image and data analysis workflow on the known lysosome positioning phenotype elicited by depletion of the small GTPase ADP-ribosylation factor-like 8b (Arl8b). It was previously shown that Arl8b loss of function leads to clustering of lysosomes close to the nucleus (Korolchuk, Saiki et al. 2011, Rosa-Ferreira and Munro 2011). Consistently, we observed perinuclear clustering of lysosomes in HeLa cells depleted of endogenous Arl8b by specific siRNA (**Figure 3A, B**). Using the OrgaMapper workflow we could faithfully segment individual HeLa cells and their nuclei and detect individual lysosomes within their cytoplasm (**Figure 3A, B**). The image and data analysis also enabled assessment of general cellular parameters such as cell area, diameter, number of detected lysosomes, and the average intensity of the organelle channel within the cytoplasm (**Figure 3C-F**). Based on the lysosome detections the average gray value at the detection peak was plotted (**Figure 3G**). The measurement yielded a highly significant reduction in the average lysosomal distance with respect to the nucleus in Arl8b-depleted cells, as expected (**Figure 3H)**. This distance measurement can be normalized to the measured Feret’s diameter of each cell, thereby reducing any measurement error due to variability in cell size (**Figure 3I**). Furthermore, the provided analysis enabled the visualization of the localization of organelles by plotting the distribution of detected organelles with respect to the nucleus as a kernel density plot. This analysis showed in more detail that the reduction in the average distance of lysosomes is an accumulation of lysosomes close to the nucleus **(Figure 3J, K**). Together, these results validate the OrgaMapper workflow as a platform capable of extracting comprehensive parameters from microscopy images and faithfully detecting alterations in lysosome positioning.

**Fig. 3:**
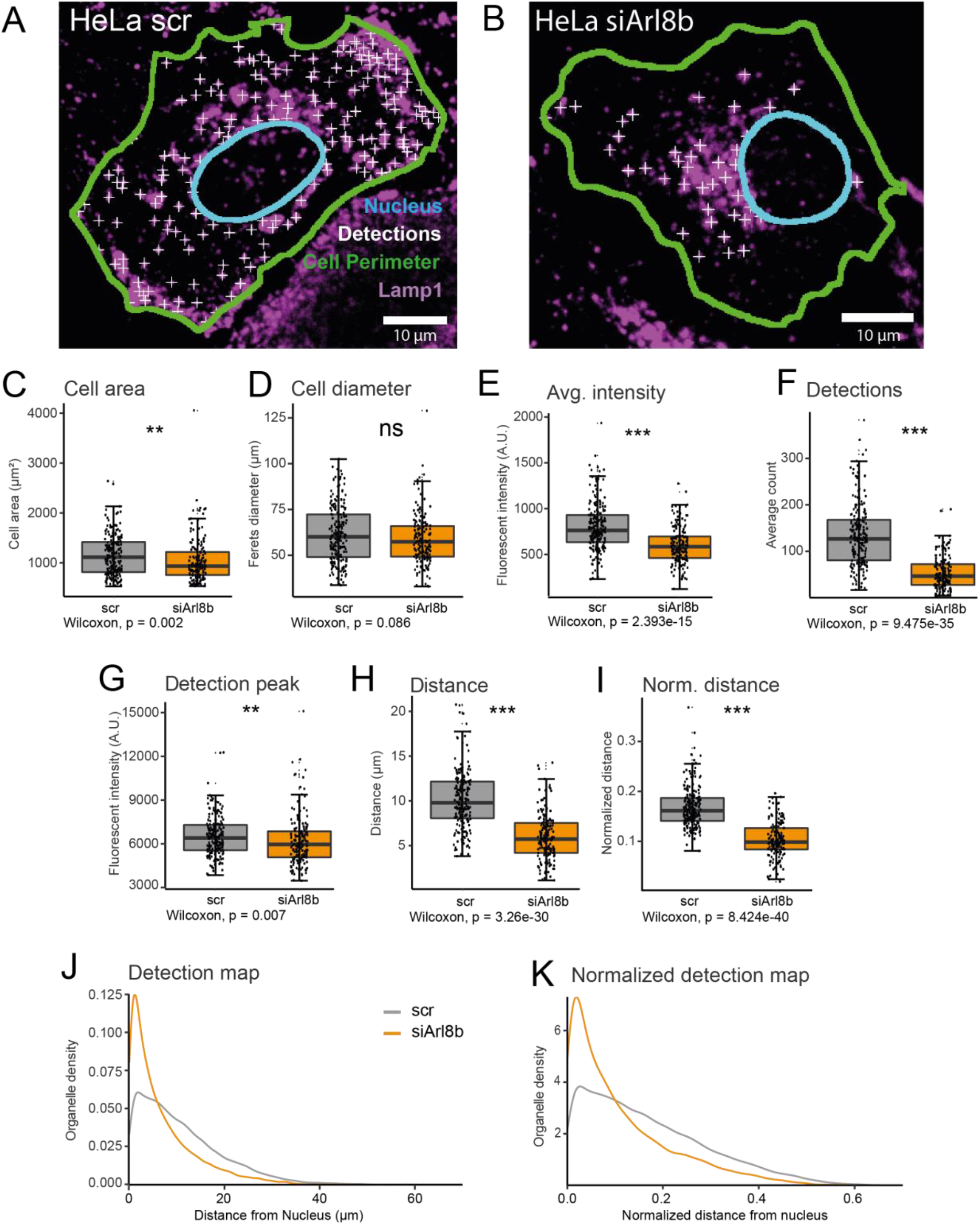
Lysosome positioning analysis with OrgaMapper. (**A**) Lysosome staining overlaid with the outline of segmented nuclei and segmented cells (green line) in HeLa cells transfected with scrambled control siRNA (scr) (**B**) or siRNA targeted against Arl8b (siArl8b). Lysosome detections are indicated by white crosses. OrgaMapper analysis extracts parameters such as (**C**) cell area, (**D**) cell diameter, (**E**) average intensity of the organelle channel, (**F**) number of detections, and (**G**) organelle intensity at the detection peak. (**H, I**) The analysis shows a significant reduction in the raw distance as well as normalized distance from the nucleus in Arl8b-knockdown cells. (**J, K**) Organelle density plots to visualize an increase in perinuclear lysosomes in knockdown cells in detail.

### OrgaMapper controls for altered cell size and organelle morphology

OrgaMapper could faithfully detect changes in lysosome positioning, a cellular process that is intimately linked to cell growth (Korolchuk, Saiki et al. 2011). We therefore wanted to validate whether the distance-based OrgaMapper analysis is indeed robust against changes in cell sizes. To this end, we utilized C2C12 wild-type and MTM1 knockout cells (Samso, Koch et al. 2022). MTM1 knockout causes a drastic reduction in cell size but no apparent phenotype in lysosome positioning (**Figure 4A, B**). As expected, OrgaMapper analysis yielded a significant decrease in cell area **(Figure 4C)** and cell diameter **(Figure 4D)** in MTM1 knockout cells. Analysis of the raw lysosome distance without cell size normalization suggested a significant reduction in the lysosome distance upon MTM1 knockout (**Figure 4E**), contrary to the visual impression. Consistent with the summary statistics, plotting the detailed lysosomal distribution indicated a slight redistribution of lysosomes closer to the nucleus (**Figure 4G**). However, normalizing lysosome distance to cell diameter abrogated the difference between wild type and MTM1 knockout cells in lysosome distance measurement (**Figure 4F**), and in the normalized distance map (**Figure 4H**). These results indicate that disregarding cell morphology can give rise to false positives and demonstrate that the OrgaMapper analysis workflow can correct for such errors.

**Fig. 4:**
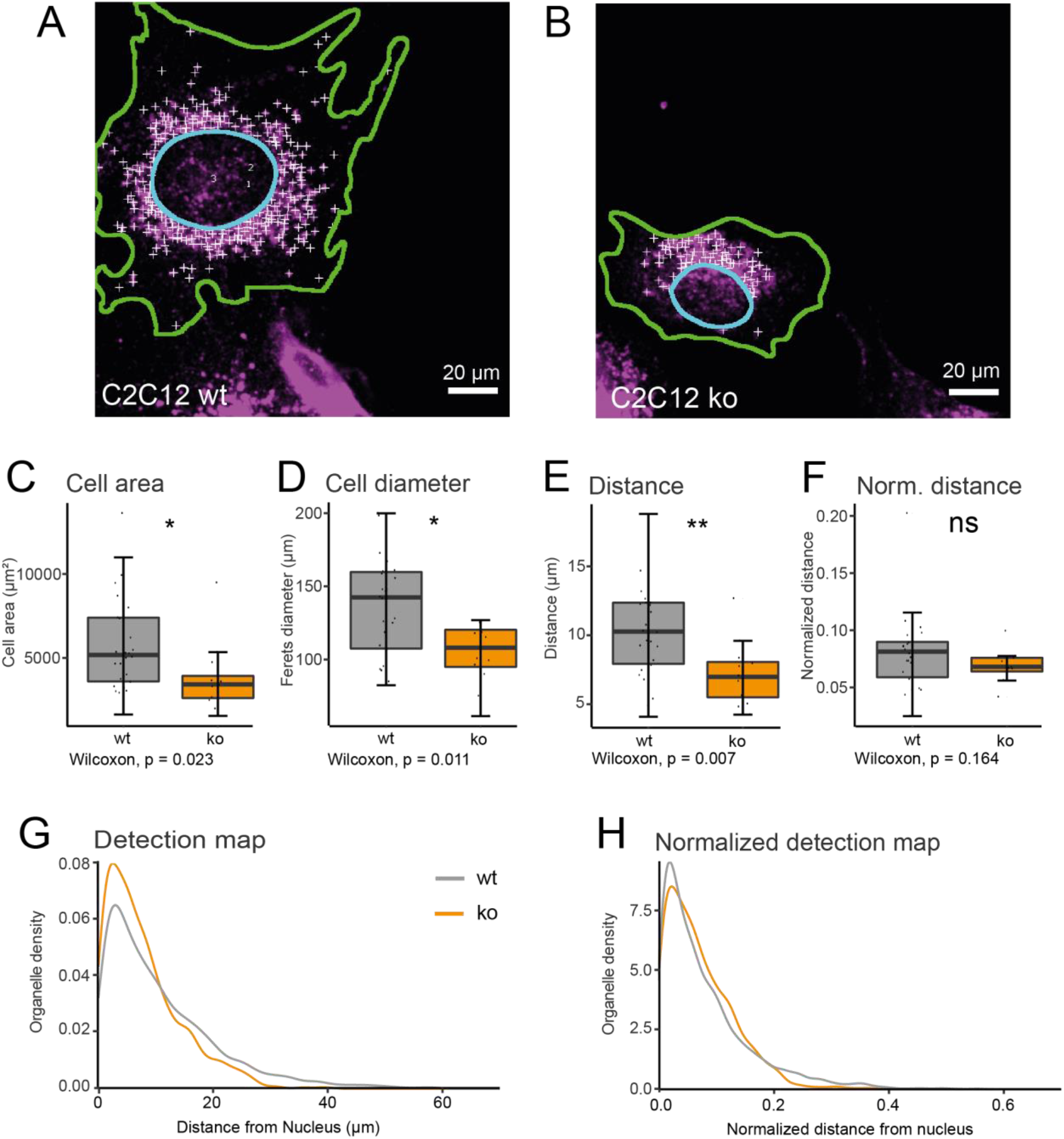
OrgaMapper analysis is robust to changes in cell size. (**A**) Lysosome staining overlaid with an outline of nucleus (cyan line) and cell segmentation (green line) in C2C12 wild type (wt) and (**B**) C2C12 MTM1 knockout (ko) cells. Lysosome detections are indicated by white crosses. (**C, D**) MTM1 knockout significantly reduces cell area and cell diameter. (**E**) Raw distance measurements indicate a significant reduction of lysosome distance to the nucleus in MTM1 knockout cells. (**F**) The apparent lysosome positioning change is abolished upon normalization of the data to cell diameter. (**G, H**) Distance maps reveal a lack of difference in lysosome localization when distance measurements are normalized to cell size.

Another source of error could arise from altered organelle morphology **(Figure 1J-L)**. In order to test whether OrgaMapper could account for such an error source we used inhibition of the phosphoinositide kinase PIKfyve by the specific inhibitor Apilimod. Apilimod treatment leads to swelling of EEA1-positive endosomes but causes no apparent endosome repositioning **(Suppl Figure 1)**. OrgaMapper could faithfully detect control and swollen endosomes and found similar distance distributions in both conditions **(Suppl. Figure 1)**. Together, these results demonstrate that OrgaMapper presents a versatile tool that can compensate for several pitfalls associated with organelle positioning measurements.

### OrgaMapper can map different organelle types

Next, we tested whether the OrgaMapper workflow is generalizable and able to assess phenotypes with respect to organelles other than lysosomes. The Golgi apparatus localizes to the perinuclear area via the microtubule organizing center. Microtubule filament disruption by Nocodazole disperses Golgi fragments into the cytoplasm **(Figure 5A, B)**. OrgaMapper could readily detect the difference in Golgi distribution between control and Nocodazole treated conditions **(Figure 5C, D)**. The dynamics and distribution of mitochondria is dependent on their anchoring to the microtubule cytoskeleton. Consistently, Nocodazole treatment causes mitochondria to cluster in the perinuclear area **(Figure 5E, F)**. In order to analyze mitochondria OrgaMapper, in addition to the spot detection, offers plots of intensity ratios or intensity maps. These faithfully detected the Nocodazole-induced perinuclear clustering of mitochondria (**Figure 5G, H**). These collective results demonstrate that the OrgaMapper workflow presents a powerful tool to map the distribution of various organelles.

**Fig. 5:**
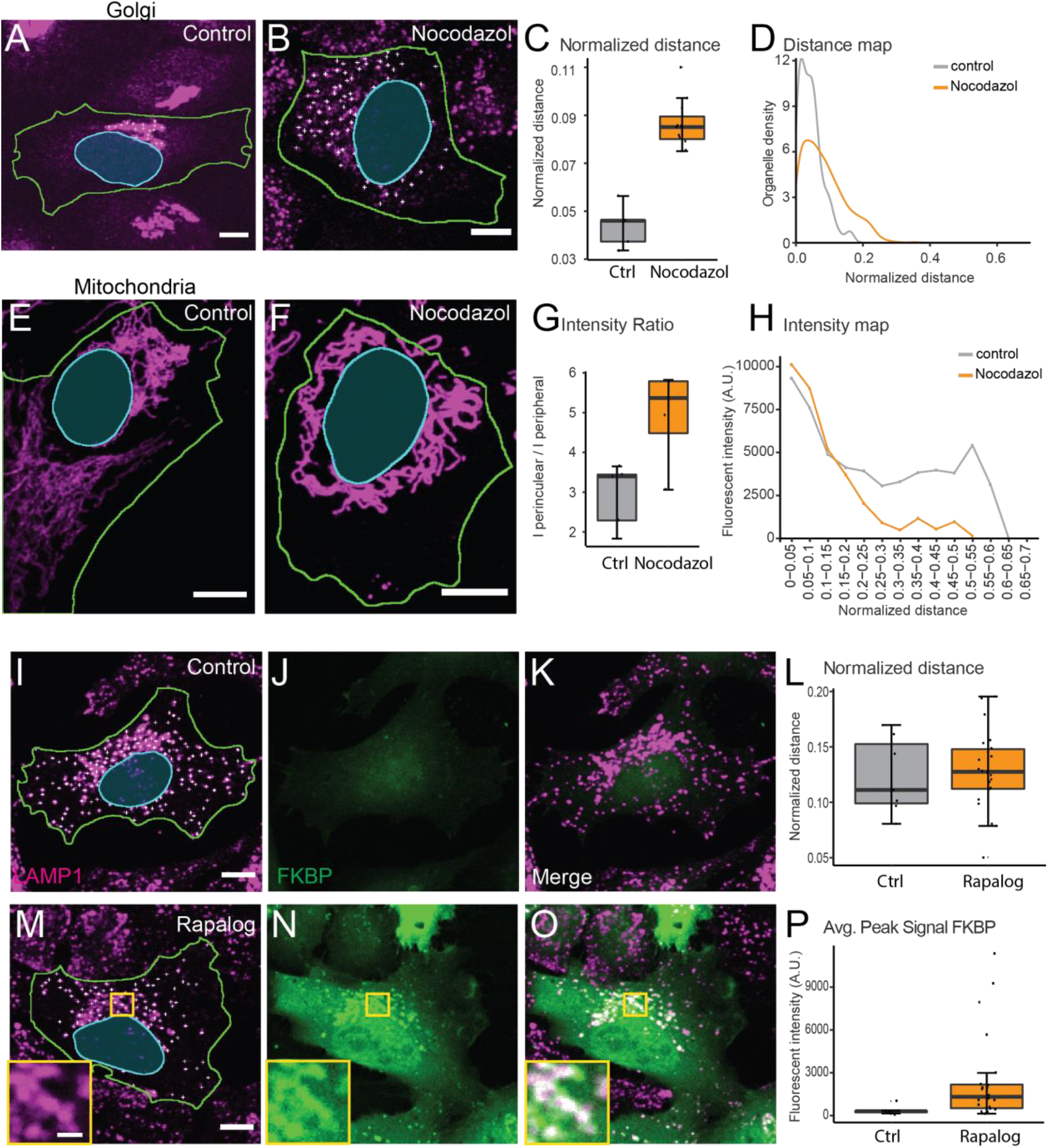
The OrgaMapper analysis can be applied to many different types of organelles. (**A-D**) OrgaMapper analysis performed on control and dispersed Golgi. (**E-H**) When detections are not possible, such as for Mitochondria, OrgaMapper offers to plot the intensity ratio as well as the intensity distribution from the nucleus. (**I-P**) OrgaMapper can measure the peak intensity as well as the intensity distribution in an additional channel. (**I-K**) Control cells expressing TMEM192-FRB (dark) and fluorescent protein tagged FKBP (**J**) do not show enrichment of FKBP on LAMP1-positive structures (**L**). (**M-O**) Treatment with Rapalog recruits FKBP to LAMP1-positive structures (**O**) which can be measured with OrgaMapper (**P**).

Organelle function and plasticity are dependent on dynamic re-localization of protein factors between the cytoplasm and the organelle membrane. For instance, dynamic Arl8b localization to lysosomes affects their intracellular distribution **(Figure 3)**. It is therefore important to examine protein abundance and localization in conjunction with organelle distribution. To enable multiplexed measurements, we extended the analysis modalities of OrgaMapper for intensity measurements in an additional channel. As a proof of principle, we utilized chemically-induced heterodimerization to enrich mRFP-FKBP on lysosomes using an FRB tagged version of the lysosomal protein TMEM192 and Rapalog, which dimerizes FKBP and FRB **(Figure 5I-K, M-O)**. Recruitment of mRFP-FKBP to lysosomes by Rapalog increased mRFP intensity on lysosomes without affecting lysosome distribution **(Figure 5L-P)**.

Together, these results demonstrate that the multimodal OrgaMapper architecture enables the assessment of the distribution of multiple organelles and the measurement of organelle-associated factors.

## Discussion

In recent years, positioning of cellular organelles in the cytoplasm has garnered substantial research interest from cell biologists and also clinicians. Each individual organelle employs a host of factors to ensure its correct positioning and the adaptation thereof should circumstances demand plasticity. However, the molecular mechanisms controlling organelle positioning dynamics remain incompletely understood. To facilitate discovery of such pathways and their molecular components robust, faithful, and scalable analysis pipelines are key. Moreover, for researchers and clinicians who choose organelle positioning as experimental or diagnostic readout easy-to-use and reliable tools are much needed. Here, we introduce OrgaMapper, an organelle positioning image and data analysis tool that is freely available, easy-to-use without in-depth image analysis knowledge, and requires freeware only, that is, Fiji and R. Moreover, OrgaMapper generates informative graphs for all measured parameters and the various means of mapping organelle and intensity distributions.

OrgaMapper is capable of analyzing organelle positioning in microscopy images based on intensity distribution of an organelle marker or based on detection of individual organelles. We tested these two approaches for robustness and faithfulness in reporting the distribution of lysosomes. We found that organelle positioning is surprisingly non-trivial to measure and, dependent on the method, can easily give rise to type I and type II errors. Using model cells and real-world specimen we show that detection of individual organelles is more reliable as compared to intensity-based approaches **(Figure 1)**. However, the detection algorithm implemented in OrgaMapper is customized for detecting vesicular structures and may not perform well in detecting organelles with tubular or sheet-like structures such as the mitochondria or the endoplasmic reticulum. It is therefore advisable to use intensity-based organelle mapping for non-vesicular organelles. Other limitations of the implemented spot detection algorithm are that vesicles with vastly different sizes or staining intensities may not be detected equally well. Likewise, sparse vesicles may be detected more faithfully than heavily crowded ones. To inform the user about the detection quality we implemented the detection preview in the Fiji plugin **(Figure 2E)**. Reviewing the detection quality should help the user to decide on whether a detection approach is feasible or whether resorting to intensity-based quantification of organelle positioning is required. To enable the detection of non-vesicular organelles future versions of OrgaMapper may include particle detection algorithms based on sophisticated automated intensity thresholds or pre-trained AI models.

We found that organelle positioning measurements are particularly sensitive to cell size. It is well known that mechanisms controlling the positioning of lysosomes are entangled with cell growth signaling pathways (Korolchuk, Saiki et al. 2011). It is therefore advisable to normalize lysosome positioning data by cell diameter, which is, however, not possible with intensity-based quantification strategies such as the widely used perinuclear index. Faithful measurement of the cell diameter relies on faithful detection of the cell edges. A prerequisite for the implementation of OrgaMapper is therefore good quality staining and image quality that is consistent among all compared samples. Future versions of OrgaMapper may implement deep learning models for single cell segmentation (Schmidt, Weigert et al. 2018, Stringer, Wang et al. 2021). Applying such a tool may abrogate the need for whole cell stains or nuclei and could free up acquisition channels for other uses such as multiple organelle stains.

In sum, we introduce a novel, easy-to-use, versatile, and robust organelle positioning image and data analysis workflow dubbed OrgaMapper. We validated the workflow using different cellular and organelle models and tested different quantification strategies. In particular, we demonstrated that measuring the distance of individually detected organelles from the nucleus edge is robust to changes in cell size, a common side effect when altering organelle positioning. OrgaMapper can be easily applied in low to high throughput microscopy-based screens to identify regulators of organelle positioning and is well suited as a validation approach for high content screening assays such as cell painting (Bray, Singh et al. 2016).

## Methods

### Organelle positioning simulation

The robustness of different organelle positioning quantification methods was tested in Fiji using *in silico* generated 3-channel images containing a single cell comprised of nucleus, cytoplasm, and organelles with defined morphology and organelle positioning. We then made incremental changes to only a single parameter (e.g. cell size) and tested how the parameter affects the result of the organelle positioning quantification. Organelle positioning was measured in the simulated cells by two methods: 1) The ratio of integrated density outside of a perimeter with defined distance to the nucleus over whole cell organelle intensity (i.e. intensity-based) and 2) the average distance of maxima detections to the nucleus (i.e. distance-based). An error factor was computed that relates a given cell parameter (i.e. cell size) to the sensitivity of the measurement results on the parameter. The error factor is computed as fold change of a given result relative to the result obtained in the cell generated with the starting parameter. For starting parameters the error factor was set to 0, i.e. the larger the error factor the more sensitive the measurement is to a given parameter.

### Cell culture

HeLa and U2OS cells were purchased from ATCC. All cell lines were maintained in DMEM containing 4.5g/ml Glucose and L-glutamine (Gibco) supplemented with 10% heat inactivated fetal calve serum (Gibco) and 100 U/ml penicillin and 100 μg/ml streptomycin (Gibco). Cells were routinely tested for Mycoplasma contamination and all tests were negative. MTM1 knockout C2C12 cell generation was described recently (Samso et al., 2022). C2C12 cells were incubated in Ringer’s solution pH 7.4 for 2 hours before fixation.

### Molecular cloning and expression constructs

For TMEM192-FRB, human TMEM192 and FKBP-Rapamycin-binding (FRB) domain of human mechanistic target of Rapamycin (mTOR) were amplified by PCR and subcloned into pEGFP/N3 vector from Clonetech by replacing EGFP.

For mRFP-FKBP, mRFP and human FK506-binding-protein 12 (FKBP) were subcloned into pcDNA3.1(+).

### Transfection and gene knockdown

Expression plasmids were transfected into cells using Lipofectamine2000 according to the manufacturer’s instructions. For siRNA transfection, 2.5×x10^5-3×x10^5 cells were seeded into 6-well dishes, reverse transfected with 50nM Arl8b-directed siRNA (GAUAGAAGCUUCCCGAAAU) or scrambled control and 6µl INTERFERin prediluted in serum-free medium, again transfected forward on day two and analyzed on day four.

### Immunofluorescence

Cells were seeded on 24 mm coverslips, cultured overnight, treated with indicated reagents, incubated with 5 µM cell tracker green CMFDA for 1 hour, fixed 15 minutes at room temperature with 4% PFA/4% sucrose in PBS, washed 3x 5 minutes with PBS, incubated with blocking buffer 1 hour room temperature, incubated with overnight at 4°C with primary antibodies diluted in blocking buffer (see below), washed 3x 5 minutes with PBS, incubated 1 hour room temperature with fluorophore-conjugated secondary antibodies in 10% goat serum/0.05% Saponin in PBS plus DAPI, washed 3x 5 minutes with PBS, mounted on ImmuMount (Shandon), and cured at room temperature.

Blocking buffer for LAMP1 (Lysosomes), EEA1 (Early Endosomes), and GM130 (Golgi) staining was 0.05% Saponin and 10% goat serum in PBS; for TOM20 (Mitochondria) blocking buffer was 0.1% TritonX-100 and 10% goat serum in PBS.

### Microscopy

All images were acquired on NikonCSU spinning disc with 40x air objective (NA = 0.95) using hardware autofocus and multi-position image acquisition. ^

### Image Analysis

Implemented as an ImageJ2 plugin in Java (Rueden, Schindelin et al. 2017) and provided via a Fiji update site. The image analysis is separated into three core tasks: segmentation of nuclei, segmentation of cell area and detection of organelles. The settings of the image analysis need to be fine-tuned for each individual imaging experiment, thus different settings for each individual image analysis component may apply. To ease the usage of OrgaMapper and to promote reproducibility the settings are saved by OrgaMapper into a XML file for each analysis run. This settings file can easily be opened again in OrgaMapper to apply the same settings with minimal effort. For the datasets in this publication the precise settings are provided for download (https://doi.org/10.5281/zenodo.8128670).

#### Nuclei segmentation

The nuclei segmentation is performed on the DAPI channel. First a median filter was applied to level out in-homogeneities in the nuclei signal without smoothing of the nuclei edge. For background subtraction a rolling ball background subtraction was applied. To segment the nuclei, an automatic global intensity threshold was applied. Optionally the segmentation mask can be adjusted using binary erosions. The particle analyzer is used to reject nuclei at the edge of the field of view as well as apply an optional size and circularity filter (**Supplementary Figure 2A**).

#### Cell segmentation

For cell segmentation the CMFDA channel was filtered using a median filter. A rolling ball background subtraction was applied to the filtered image. A fixed global intensity threshold was applied to generate binary masks of the cell area. To separate touching cells a marker-controlled watershed was used. First the signal of the nuclei channel and the CMFDA channel was added together. The composite image was then filtered using a large Gaussian blur. To determine the separation of the touching cells the find maxima algorithm was applied using the segmented particles option. This applies a watershed algorithm based on the intensity values of the combined and smoothed Nuclei and CMFDA channel and results in a binary mask containing the boundaries of touching cells. The cell area mask and the cell boundary mask were multiplied to generate a binary mask with individual cells separated. The cells were further filtered for size and circularity using the particle analyzer option in ImageJ. Further the cells were filtered if they did not contain a nuclei segmentation or more than one nuclei segmentation (**Supplementary Figure 2B**).

#### Organelle detection

To detect individual blob shaped organelles the organelle channel was filtered using an ImageJ implementation of the Laplacian-of-Gaussian filter (Sage, Neumann et al. 2005). Individual organelles were then detected using a maxima detection. The detections were filtered for excluding detections in the nuclei mask (**Supplementary Figure 2C**).

#### Measurements

We extracted key measurements per well such as total cell count and mean intensity of the background based on the area outside of the cell segmentation. For each cell we further extracted parameters such as cell area, ferret diameter and mean intensity of the organelle channel as well as an optional measurement channel within the cytoplasmic area (cytoplasmic area: cell mask minus nuclei mask). To determine the distance from the nucleus of each organelle detection an Euclidean distance map (EDM) was computed per cell. Which is a very fast computation as compared to the algorithm used by (Klickstein, Mukkavalli et al. 2020). For each individual detection within each cell the distance based on the EDM was extracted (**Supplementary Figure 2D**). Further the signal intensity at that location of the detection was measured in the organelle channel as well as an optional measurement channel. As an alternative detection independent measurement, the distance of each individual pixel within the cytoplasmic mask of each cell was extracted as well as the corresponding intensity value in the organelle channel and measurement channel.

### Statistical Analysis and data visualization

The results of the image analysis with the Fiji plugin were collected using the OrgaMapper Shiny app. Further the Shiny app allows basic and advanced descriptive data analysis. Intensity measurements were background subtracted using the background intensity measured outside of the segmented cell area in the organelle and additional measurement channel. For the analysis a cell diameter can be applied (Set to 600 µm). Distance measurements were normalized per cell using the extracted Feret’s diameter of the cell. For visualizing the organelle distribution based on the organelle detection a kernel density plot was computed. The intensity distribution was based on binning the intensity values and visualizing as a line plot. The intensity ratio was computed using a separate R script by dividing the cell in two areas using a fixed perimeter of 10 µm. The mean intensity from the area closer to the nucleus was divided by the mean intensity away from the nucleus. All measured and evaluated cellular and organelle parameters were visualized as box plots overlaid with a dot plot. The statistical significance between control and treatment was evaluated using an unpaired two-sample Wilcoxon rank sum test.

## Materials

### Reagents

| name | company | Product number | working concentration |
| --- | --- | --- | --- |
| **cell tracker green CMFDA** | Invitrogen | C7025 | 5 μM |
| **Nocodazole** | Sigma | M1404 | 20 μM |
| **Apilimod** | Sigma | SML2974 | 1 μM |
| **A/C Heterodimerizer (Rapalog)** | Takara | 635095 | 500 nM |
| **Arl8b siRNA** | Sigma |  | 100 nM |
| **Sigma Mission® siRNA universal negative control #1** | Sigma | SIC001 | 100 nM |

### Antibodies

| name | company | Product number | dilution |
| --- | --- | --- | --- |
| **anti-LAMP1** | BD pharmingen | 555798 | 1:200 |
| **anti-TOM20** | Santa Cruz | sc-17764 | 1:200 |
| **anti-GM130** | Abcam | ab52649 | 1:200 |
| **anti-EEA1** | Cell Signaling | 2411 | 1:100 |
| **Anti-rabbit CF647** | Biotium | 20047 | 1:1000 |
| **Anti-mouse CF647** | Biotium | 20046 | 1:1000 |

