## Supplemental Figures for "OrgaMapper: A robust and easy-to-use workflow for analyzing organelle positioning"

### Supplemental Data & Figures

#### Supplemental Figure 1

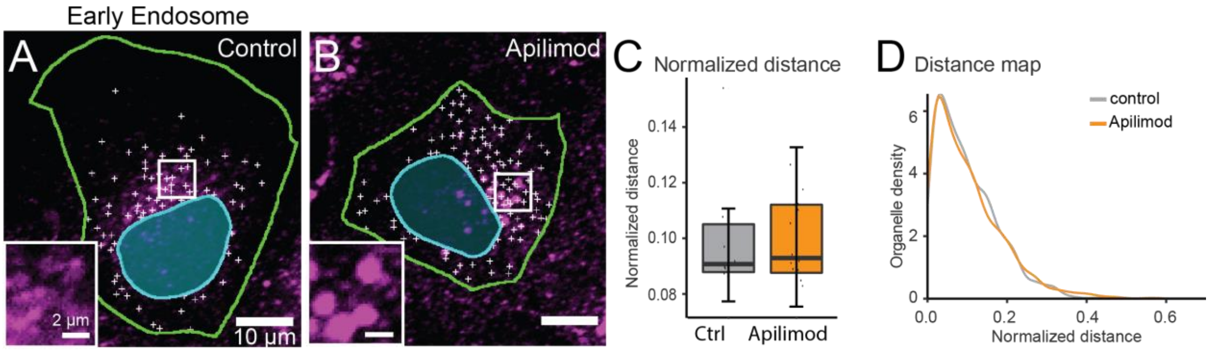


**Suppl. Fig. 2:** OrgaMapper analysis performed on early endosomes marked by an EEA1-specific antibody in control (**A**) and Apilimod treated conditions in which endosomes undergo swelling (**B**). Normalized distance of detected endosomes to the nucleus (**C**) and distance mapping (**D**) do not show a difference in organelle positioning.

Supplemental Figure 2

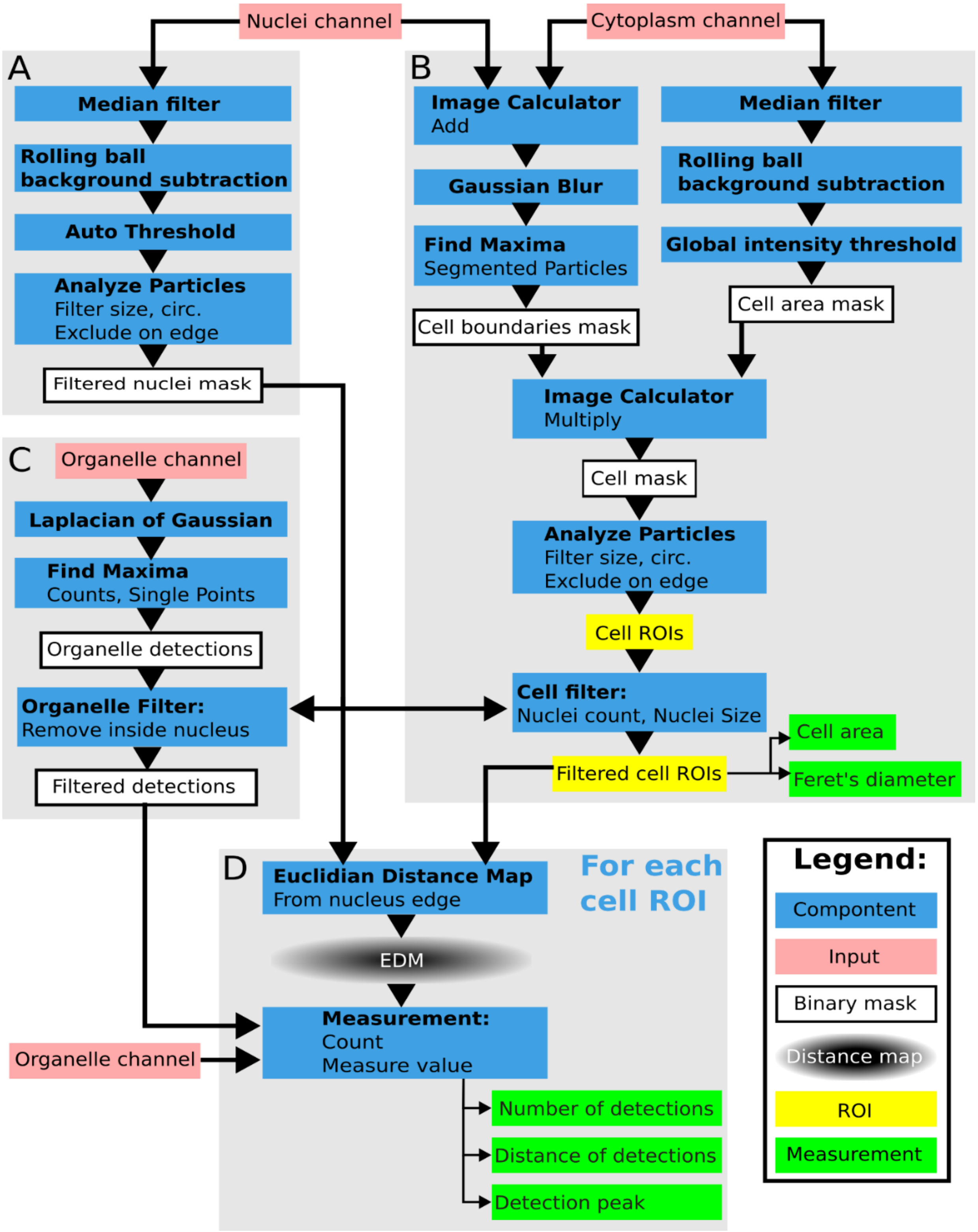


**Suppl. Fig. 2: OrgaMapper workflow diagram of main modules.** (**A**) Nuclei segmentation. (**B**) Cell segmentation. (**C**) Organelle detection. (**D**) Organelle measurement.
